# An impressive capacity for cold tolerance plasticity protects against ionoregulatory collapse in the disease vector, *Aedes aegypti*

**DOI:** 10.1101/745885

**Authors:** Amanda Jass, Gil Y. Yerushalmi, Hannah E. Davis, Andrew Donini, Heath A. MacMillan

**Affiliations:** Department of Biology, York University, Toronto, Canada, M3J 1P3; Department of Biology, Carleton University, Ottawa, Canada, K1S 5B6

**Keywords:** Chill tolerance, disease vector, hyperkalemia, ion balance, phenotypic plasticity, thermal performance

## Abstract

1. The mosquito *Aedes aegypti* is largely confined to tropical and subtropical regions but its range has recently been spreading to colder climates. As insect biogeography is closely tied to environmental temperature, understanding the limits of *Ae. aegypti* thermal tolerance and their capacity for phenotypic plasticity is important in predicting the spread of this species.
2. In this study we report on the chill coma onset and recovery, as well as low temperature survival phenotypes of larvae and adults of *Aedes aegypti* that developed or were acclimated to 15°C (cold) or 25°C (warm).
3. Developmental cold acclimation did not affect chill coma onset of larvae but substantially reduced chill coma onset temperatures in adults. Chill coma recovery time was affected by both temperature and the duration of exposure, and developmental and adult acclimation both strongly mitigated these effects and increased rates of survival following prolonged chilling.
4. Female adults were far less likely to take a blood meal when cold acclimated and simply exposing females to blood (without feeding) attenuated some of the beneficial effects of cold acclimation on chill coma recovery time.
5. Lastly, larvae suffered from hemolymph hyperkalemia when chilled, but development in the cold attenuated the imbalance, which suggests that acclimation can prevent cold-induced ionoregulatory collapse in this species.
6. Our results demonstrate that *Aedes aegypti* larvae and adults have the capacity to acclimate to cold temperatures and do so at least in part by better maintaining ion balance in the cold. This ability for cold acclimation may facilitate the spread of this species to higher latitudes, particularly in an era of climate change.

## Introduction

The mosquito *Aedes aegypti* is abundant in tropical and subtropical regions where it is an arboviral disease vector for Zika, Chikungunya, yellow fever, and dengue (Bhatt et al., 2013; Kraemer et al., 2019, 2015). The global distribution of *Ae. aegypti* is closely related to environmental temperatures (Brady et al., 2014, 2013; Kraemer et al., 2019). This pattern suggests that like other small dipterans such as *Drosophila*, the inability of *Aedes* to survive cold winters in poleward latitudes limits its ability to colonize these areas (J. L. Andersen et al., 2015; Kellermann et al., 2012; Overgaard, Kearney, & Hoffmann, 2014).

In recent years, however, *Ae. aegypti*, and the closely-related vector *Ae. albopictus*, have spread through the northeastern United States. Adults of both species have even begun to appear in southern Ontario, Canada early in spring (mosquito responsible for majority of Zika infections found in Canada for the first time, 2017), which suggests successful local overwintering of at least some adults or late-stage pupae. The ability to overwinter in northern climates is completely at odds with our understanding of *Ae. aegypti* as a cold-intolerant species with little to no ability for seasonal quiescence or diapause (Diniz, Albuquerque, Oliva, Melo-santos, & Ayres, 2017). Current climate models predict continued increases in average global temperatures and a greater frequency of extreme thermal events during winter (Easterling et al., 2000; Williams, Henry, & Sinclair, 2014), but predictive models of *Ae. aegypti* distribution do not currently consider the possibility of phenotypic plasticity in this species (e.g. Kamal et al., 2018), because no such plasticity has been described.

Because of their global importance as disease vectors and their demonstrated potential for invasion, there is growing interest in understanding the limits of mosquito thermal tolerance. Low temperatures adversely affect life history traits throughout the *Ae. aegypti* life cycle, including development rate, reproductive success, and survival (Carrington, Armijos, Lambrechts, Barker, & Scott, 2013; Davis, 1931; Yang, Macoris, Galvani, Andrighetti, & Wanderly, 2009). The effects of temperature on *Ae. aegypti* developmental success appear quite pronounced, as survival from egg to adult drops from 92% at 20°C to only 3% at 15°C (Rueda, Patel, Axtell, & Stinner, 1990). Of the different life stages of *Ae. aegypti*, the eggs appear tolerant to cold, surviving and hatching following cold exposures up to 24 h at −2°C or 1 h at −17°C (Davis, 1931; Thomas, Obermayr, Fischer, Kreyling, & Beierkuhnlein, 2012). Accordingly, *Ae. aegypti* have been documented to successfully overwinter as far north as Washington D.C. and Indiana, and as far south as Buenos Aires, and there is evidence of cold adaptation occurring in these temperate populations (De Majo, Montini, & Fischer, 2017; Fischer, Alem, De Majo, Campos, & Schweigmann, 2011; Hawley, Pumpuni, Brady, & Craig, 1989; Lima, Lovin, Hickner, & Severson, 2016). Like many other poikilotherms, larval *Ae. aegypti* experience slowed development, delayed and decreased pupation, and increased mortality with decreasing temperatures (Brady et al., 2014; Carrington et al., 2013; De Majo et al., 2017; Tun-Lin, Burkot, & Kay, 2000; Yang et al., 2009). Similarly, adult *Ae. aegypti* experience increased mortality, decreased oviposition rate, and overall reduced fecundity at 15°C (Tun-Lin et al., 2000). To date, however, studies of temperature effects on *Ae. aegypti* have largely focused on consequences to reproductive success, growth and development, and there has been little work focused on the extreme limits of thermal tolerance or the potential for thermal plasticity, particularly in later life stages. This gap in knowledge represents a considerable risk, particularly considering recent reports from *Drosophila* that thermal limits may better predict species distribution and abundance than optimal temperatures or rates of growth and reproduction at more favourable temperatures (MacLean et al., 2019; Overgaard et al., 2014).

*Ae. aegypti* is a chill susceptible insect, meaning it succumbs from exposure to low temperatures well above the freezing of its bodily fluids. The cold tolerance of chill susceptible insects can vary widely, both among and within species. Broad differences in basal cold tolerance can exist among populations or species (Gibert, Moreteau, Pétavy, Karan, & David, 2001; Kellermann et al., 2012; Vorhees, Gray, & Bradley, 2013; Warren & Chick, 2013), and many insects can also drastically alter their cold tolerance within their lifetime. For example, insects can undergo thermal acclimation in response to chronic low temperature exposure, or rapidly harden in response to an acute temperature change (e.g. rapid cold-hardening) (Colinet & Hoffmann, 2012; Hoffmann, Scott, Partridge, & Hallas, 2003; Kellermann et al., 2012; Kelty & Lee, 2001; Sinclair et al., 2006). To date, a capacity for thermal plasticity at low temperature (cold acclimation) has been demonstrated in many chill-susceptible insects, such as fruit flies, cockroaches, locusts, and crickets (M. K. Andersen, Folkersen, MacMillan, & Overgaard, 2017; Coello Alvarado, MacMillan, & Sinclair, 2015; Colinet & Hoffmann, 2012; V Koštál, Yanagimoto, & Bastl, 2006).

Cold acclimation typically affects a variety of cold tolerance phenotypes in chill susceptible insects. For example, cold-acclimated insects commonly have a lower temperature of chill coma onset (CCO), more rapidly recover from chill coma following rewarming (a lower chill coma recovery time; CCRT), and avoid the development of cold-induced injury better than warm-acclimated conspecifics (Coello Alvarado et al., 2015; MacMillan, Andersen, Loeschcke, & Overgaard, 2015; Ransberry, MacMillan, & Sinclair, 2011). While little is known about cold acclimation in *Ae. aegypti*, eggs of *Ae. albopictus* have increased cold tolerance following cold acclimation (Hanson & Craig, 1995). The magnitude of cold plasticity can vary among and within populations (Nyamukondiwa, Terblanche, Marshall, & Sinclair, 2011; Sørensen, Kristensen, & Overgaard, 2016). In the case of *Ae. albopictus*, cold acclimation was only noted in temperate populations and not tropical populations, so this capacity for plasticity is thought to be facilitating the northward expansion of the species’ range (Hanson & Craig, 1995; Rochlin, Ninivaggi, Hutchinson, & Farajollahi, 2013; Romi, Severlini, & Toma, 2006).

While tolerance to extreme cold relies on a physiological capacity to avoid or survive ice formation inside the body, tolerance to chilling requires a physiological capacity to resist the effects of low temperature *per se* on organ, tissue, and cellular biochemistry (MacMillan, 2019; Overgaard & MacMillan, 2017; Teets & Denlinger, 2013). Consequently, measures of cold tolerance relevant to freeze avoidant and freeze tolerant insects, such as the supercooling point (the temperature of spontaneous ice formation within the body) or survival following freezing, are irrelevant to characterizing the thermal limits of chill susceptible insects (Overgaard & MacMillan, 2017). When cooled below a critical threshold temperature, chill susceptible insects suffer a local loss of ion homeostasis in the nervous system, leading to nerve depolarization (spreading depression) and a state of complete neuromuscular silence termed chill coma (MacMillan & Sinclair, 2011a; Mellanby, 1939; Robertson, Spong, & Srithiphaphirom, 2017). The temperature at which this paralytic state occurs is called the chill coma onset temperature (CCO) (Overgaard & MacMillan, 2017). With time spent at low temperatures, chill susceptible insects lose ion and water balance and suffer from hemolymph hyperkalemia (high [K^+^]), which further depolarizes cells and activates voltage-gated calcium channels, driving rampant cellular apoptosis (Bayley et al., 2018; MacMillan, Andersen, Davies, & Overgaard, 2015; MacMillan, Baatrup, & Overgaard, 2015). The severity of this loss of homeostasis increases with longer or lower temperature exposures, and the tissue damage that accrues while an insect is in this state is thought to largely determine its survival and fitness following rewarming (Overgaard & MacMillan, 2017). Species that are more cold tolerant, or individuals that have acclimated to low temperatures are better able to maintain ion and water balance during cold exposure (M. K. Andersen, Folkersen, et al., 2017; Coello Alvarado et al., 2015; V Koštál et al., 2006; MacMillan, Andersen, Loeschcke, et al., 2015).

Here, we use a laboratory-bred population of *Ae. aegypti* to determine chill coma onset and recovery phenotypes of larvae and adults of this species. We allowed larval and adult mosquitoes to undergo either warm (25°C) or cold acclimation (15°C) to test whether this species is capable of acclimating to sub-optimal thermal conditions. Cold acclimation led to significant changes in the cold tolerance of both larvae and adults, so we used larvae to test whether improvements in cold tolerance following cold acclimation are driven by an improved ability to maintain ion balance in the cold.

## Materials and Methods

### Animal husbandry

A colony of *Aedes aegypti* mosquitoes (Linnaeus) was established in 2007 at York University from eggs provided by M. Patrick (San Diego) and supplemented with eggs from Liverpool strain provided by C. Lowenberger (Simon Frasor, BC., Canada). Our mosquitoes were reared as described by Misyura et al. (2017) with slight modifications. Briefly, eggs were hatched in 2 L of dechlorinated tap water (water changed every 4 d) and fed 6 mL of a premade food solution composed of 1.8 g liver powder and 1.8 g of inactive yeast in 500 mL of reverse-osmosis water daily. The population is maintained at room temperature (22±1°C) with a 12:12 h light:dark cycle.

To obtain larvae for experiments, eggs were added to 1.5 L of dechlorinated tap water along with 2 mL of the liver-yeast diet. Containers of water and eggs were kept in a 25 ± 0.5°C incubator (12 h:12 h light:dark). The next day, hatching was confirmed through visual inspection and 2 mL of food was added. One day later, all larvae in each bin were randomly assigned to one of two developmental acclimation treatments, 15°C or 25°C, such that each treatment had approximately equal numbers. The larvae assigned to each treatment were transferred to a new container filled with 1.5 L dechlorinated tap water and 2 mL food and placed in either the 15°C or the 25°C incubator. Larvae were fed 2 mL liver-yeast food mix every day until the first pupa was spotted. Water was changed as needed, with food always added after a water change. When the first pupae were observed, 4^th^ instar larvae were collected to be used in experiments.

To acclimate adults for experiments, pupae (reared under standard colony conditions as described above) were isolated daily and given 1-2 days to mature prior to the placement of 40 ± 10 pupae in small open-top containers with ~60 mL of dechlorinated tap water. The open-top containers were then placed within custom made (18 cm long × 15 cm wide × 10 cm tall) enclosed containers (with a netted section to allow for air flow) allowing the pupae to emerge over a period of 48 h. A premade sugar water solution (40 g of sucrose in 250 mL of tap water) was placed in each container to allow for adults to feed. Following 48 h given for emergence, any remaining pupae were removed, and the containers were separated into two different acclimation treatments: cold-acclimation (15°C) and warm-acclimation (25°C). This ensured all adults were 1-2 days old upon the initiation of the acclimation treatments. Both acclimation groups were maintained on a 12:12 h light:dark cycle. Adult mosquitoes were left at their respective acclimation temperatures for five days, and thus all adults were 6-7 days post-emergence when used in experiments.

### Chill coma onset

To assess chill coma onset temperatures (CCO), individual larvae were collected from the developmental acclimation treatments using a pipette and transferred to 4 mL glass vials along with 2 mL of their rearing water. The vials were affixed to a custom-made aluminum rack that was submerged in a glass aquarium containing a 1:1 mixture of ethylene glycol and water, which was circulated by a programmable refrigerated bath (Model AP28R-30, VWR International, Mississauga, ON, Canada). The temperature of the bath was independently monitored with a pair of type-K thermocouples connected to a computer running Picolog (version 5.25.3) via a Pico TC-07 interface (Pico Technology, St. Neots, UK). The larvae were held at 20°C for 15 min then ramped down at 0.1°C/min. We recorded the temperature at which each larva completely stopped responding to vibrational and light stimuli. Since larvae would often ignore a stimulus during one scan only to respond strongly on the next, all larvae, including those that had been recorded as being in chill coma, were tested for a response throughout the experiment to ensure the accuracy of the CCO temperature.

Adult mosquitoes from the 25°C and 15°C acclimation groups were briefly anesthetized under CO_2_ and placed in 4 mL glass vials (filled with ambient air) and affixed to a rack that was submerged in a temperature-controlled bath, as described above for the larvae. To record adult CCO, the temperature of the bath was initially set to 25°C for 15 min and then ramped down at a rate of 0.13°C min^−1^ while mosquito movement was continuously monitored. The temperature at which movement stopped following perturbation with a plastic probe was recorded as the adult mosquito CCO.

### Chill coma recovery time (CCRT)

To measure chill coma recovery time (CCRT), larvae were exposed to 2°C for 4, 8, 12, or 16 hours. Larvae were cold-exposed by transferring each individual to a 1.5 mL open centrifuge tube and incubating the tubes in a refrigerated centrifuge (Thermo ScientificTM Sorvall LegendTM Micro 21R) set to 2°C (temperature was confirmed via independent thermocouples, and chosen based on prior trials). After exposure to the cold, each larva was transferred to its own 6.7cm diameter plastic container, filled with 50 mL room temperature dechlorinated water. A timer was set immediately upon placement of the larva into room temperature water. CCRT was assessed as the time taken for the larva to swim a continuous distance of 2 cm (measured using a 1 cm^2^ grid lining the bottom of the container). Larvae that could not swim 2 cm within 2 h were considered to have suffered severe injury.

Chill coma recovery time (CCRT) was determined in adult mosquitoes following 6 h at 2°C. Mosquitoes were sexed and placed in 4 mL enclosed glass containers at room temperature (22±1°C) and observed for 120 minutes. The duration of time required for a mosquito to stand on all 6 legs following its removal from the cold was recorded as its CCRT. To assess the effect of blood feeding on CCRT, sugar-water mixture was removed from the cages of warm-acclimated mosquitoes 24 h before blood feeding. Mosquitoes were exposed to warm sheep blood for 20 min through a thinly stretched parafilm membrane. The mosquitoes were then either given a 0, 40, or 160 min (or alternatively 20, 60, or 180 min from the onset of blood feeding) period prior to the initiation of the cold treatment of 6 h at 2°C. Mosquitoes that did not feed during the 20 min blood exposure period were used as an internal control.

### Low temperature survival

To measure low temperature survival of the larvae, groups of 24 larvae were exposed to −4°C, −2°C, 0°C, 2°C, 5°C, or 10°C for 24 h using a refrigerated centrifuge as described for CCRT. Importantly, the water containing the larval mosquitoes was never observed to freeze under any of these conditions (i.e. the water supercooled). After exposure to the cold, larvae were kept at room temperature for an additional 24 h, and then survival proportion was recorded. Larvae that were able to move when disturbed were counted as alive. Chilling injury was measured as the inability to resume use of the siphon, where larvae that could not use their siphon to ventilate within 2 hours were counted as injured.

Chilling-survival was assessed in adult mosquitoes following 6 h exposures to temperatures between −4 and 2°C. The exposure temperatures varied somewhat between the two acclimation groups to include temperatures that result in survival proportions ranging from 0% to 100%. To this end, cold-acclimated mosquitoes were exposed to −4°C, −3°C, −2.5°C, −2°C, −1°C, 0°C, 1°C, and 2°C and warm-acclimated mosquitoes were exposed to −2°C, −1°C, −0.5°C, 0°C, 1°C, and 2°C. Immediately upon removal from the cold exposure mosquitoes were isolated in 4 mL enclosed glass containers and left at room temperature (22 ± 1°C) for 18 h to recover. Following this, the mosquitoes were assessed such that those that were able to stand were considered alive while those that were unable to stand were considered dead.

### Hemolymph ion concentration

To quantify Na^+^ and K^+^ concentrations in larval hemolymph, we used the ion-selective microelectrode technique (ISME). Control larvae were sampled directly from their rearing conditions, while cold exposed larvae were first exposed to 24h at 0°C (using a refrigerated centrifuge as described for CCRT), before hemolymph was sampled and immediately measured. Hemolymph was collected by first securing larvae onto lids from 35 mm × 10 mm sterile petri dishes using Murray’s® pure beeswax. Each larva was immobilized by applying beeswax to the head and terminal segment. A drop of paraffin oil was then applied to the abdomen, and the cuticle of this region was lightly sheared open with a sharp-pointed metal pin. The emerging droplet of hemolymph was collected and placed under mineral oil in a petri dish coated with a silicone elastomer using a micropipette.

Custom-made ion-selective microelectrodes were constructed and used following previously described methods (Jonusaite, Kelly, & Donini, 2011). Briefly, borosilicate glass capillaries were pulled to a tip diameter of ~3 μm using a micropipette puller (Flaming Brown P-97, Sutter Instruments, Novato, USA), heated to 300°C, and exposed to N,N-dimethyltrimethylsilylamine vapour for 1 h. Potassium-sensitive electrodes were backfilled with 100 mM of KCl and front-filled with K^+^ ionophore (K^+^ ionophore I, cocktail B, Sigma Aldrich, St. Louis, MO, USA). Sodium sensitive electrodes were backfilled with 100 mM NaCl and front-filled with Na^+^ ionophore (Na^+^ Ionophore II Cocktail A; Sigma Aldrich). The circuit was completed with a reference electrode pulled from filamented glass capillary and back-filled with 500 mM KCl. Signal information was relayed to a PowerLab 4/30 data acquisition device (ADInstruments; Sydney, AUS) and interpreted by LabChart 6 software (ADInstruments). Voltages obtained from the hemolymph samples were compared to those from calibration solutions of known concentrations, and the Nernst slope was applied to determine hemolymph ion concentration ([X]) using the following formula:

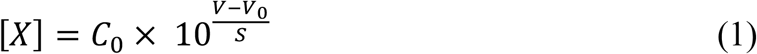

Where *C*_0_ is the lower calibration concentration in mM, *V* is the voltage (mV) reading from the hemolymph sample, *V*_0_ is the voltage (mV) reading of the lower calibration concentration, and *S* is the slope of the electrode (mV) which is the difference in voltage between the two calibration solutions that differ in concentration by a factor of 10. The following calibration solutions were used: Na^+^ (20 mM NaCl/180 mM LiCl, and 200 mM NaCl); and K^+^ (0.5 mM KCl/49.5 mM LiCl, 5 mM KCl/45 mM LiCl, and 50mM KCl).

### Data analysis

R (version 3.6.1) was used to complete all data analyses (R Development Core Team, 2019). Larval CCO temperatures were compared between acclimation groups using a one-way ANOVA and CCRT temperatures with a generalized linear model (GLM) with exposure time and acclimation temperature as factors. Rates of injury in larval mosquitoes following the CCRT assays were compared using a two-way ANOVA with acclimation temperature and duration of cold exposure included as factors. Adult CCO and CCRT were both compared using two-way ANOVAs (with acclimation temperature and sex as factors). The effect of blood feeding on CCRT in adult mosquitoes was analyzed for each acclimation group independently (because cold acclimated mosquitoes did not feed) using GLMs. Feeding status and time since feeding were included as factors for warm acclimated mosquitoes, and only time since feeding for cold-acclimated mosquitoes. Survival following cold stress was analyzed for each life stage using GLMs with a binomial error distribution and a logit-link function. Acclimation treatment and temperature were included as factors for larval survival, and sex, acclimation treatment, and temperature were included for the adults. Initial models were saturated with all potential interactions and were reduced to find the most parsimonious model based on Akaike’s Information Criterion (ΔAIC > 2).

## Results

### Chill coma onset

Developmental acclimation did not significantly affect chill coma onset (CCO) temperatures in larval mosquitoes (F_1,67_ = 1.3; *P* = 0.265); both warm- and cold-acclimated mosquitoes had a CCO of ~6°C (Fig. 1A). The majority of larvae were observed to sink to the bottom of the glass vials upon entering chill coma. Although no effect was seen in the larvae, cold acclimation strongly reduced CCO temperatures in both male and female adult mosquitoes (Fig. 1B; main effect of acclimation: F_3,27_ = 48.5, *P* < 0.001). The magnitude of this effect differed between the sexes; while the mean female CCO differed by ~3°C, that of males differed by ~6.4°C (Fig 1B; interaction between acclimation status and sex: F_3,27_ = 4.8, *P* = 0.037). In general, females tended to have lower CCO temperatures than males (main effect of sex: F_3,27_ = 6.8, *P* < 0.014).

**Figure 1:**
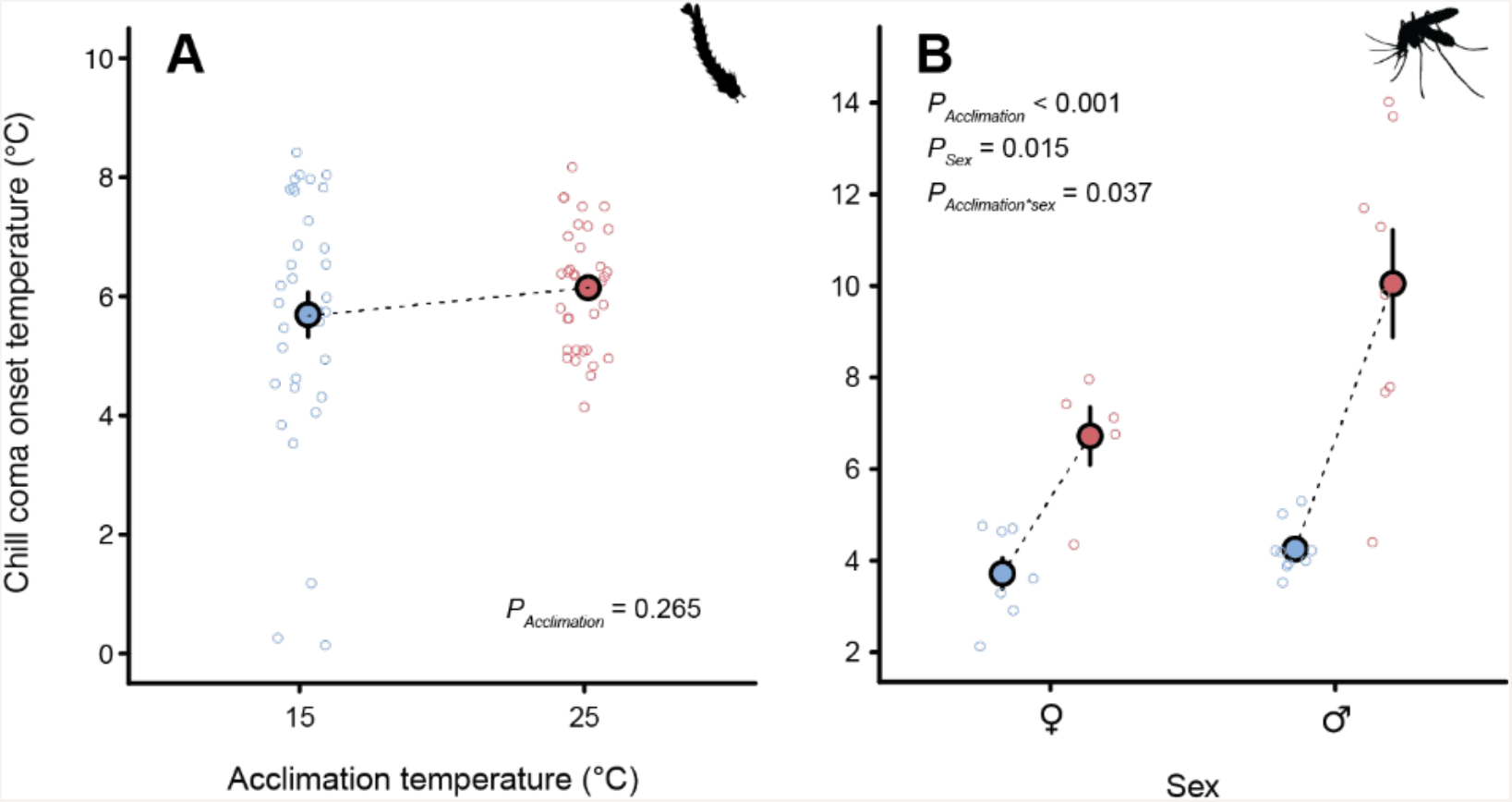
Chill coma onset temperatures of larval (A) and adult (B) *Aedes aegypti* acclimated to warm (25°C; red) and cool (15°C; blue) conditions. Open circles represent individual mosquitoes and closed circles represent the mean (± sem) for each acclimation group and life stage. Error bars that are not visible are obscured by the symbols.

### Chill coma recovery and injury

Developmental acclimation strongly impacted chill coma recovery time (CCRT) following exposure to 2°C in larval mosquitoes. Acclimation temperature and exposure duration interacted to determine CCRT (Fig. 2A; F_3, 552_ = 20.6, *P* < 0.001), such that increasing duration of cold exposure led to longer recovery times in warm-acclimated, but not cold-acclimated larval mosquitoes. In addition to increases in mean recovery times, CCRT became increasingly variable in warm-acclimated mosquitoes with increasing duration of cold stress, but the same was not true in the cold-acclimated conspecifics (Fig. 2A). The proportion of larvae that could resume use of their siphon 2 h following recovery was examined in the same individuals (an index of chilling injury). Warm-acclimated larvae suffered clear chilling injury (roughly 30% after 12 or 16 h at 2°C), while only slight chilling injury (~2%) was noted in the cold-acclimated larvae following the same exposures (Fig. 2B; interaction between acclimation and exposure duration: F_3,20_ = 10.0, *P* = 0.005).

**Figure 2:**
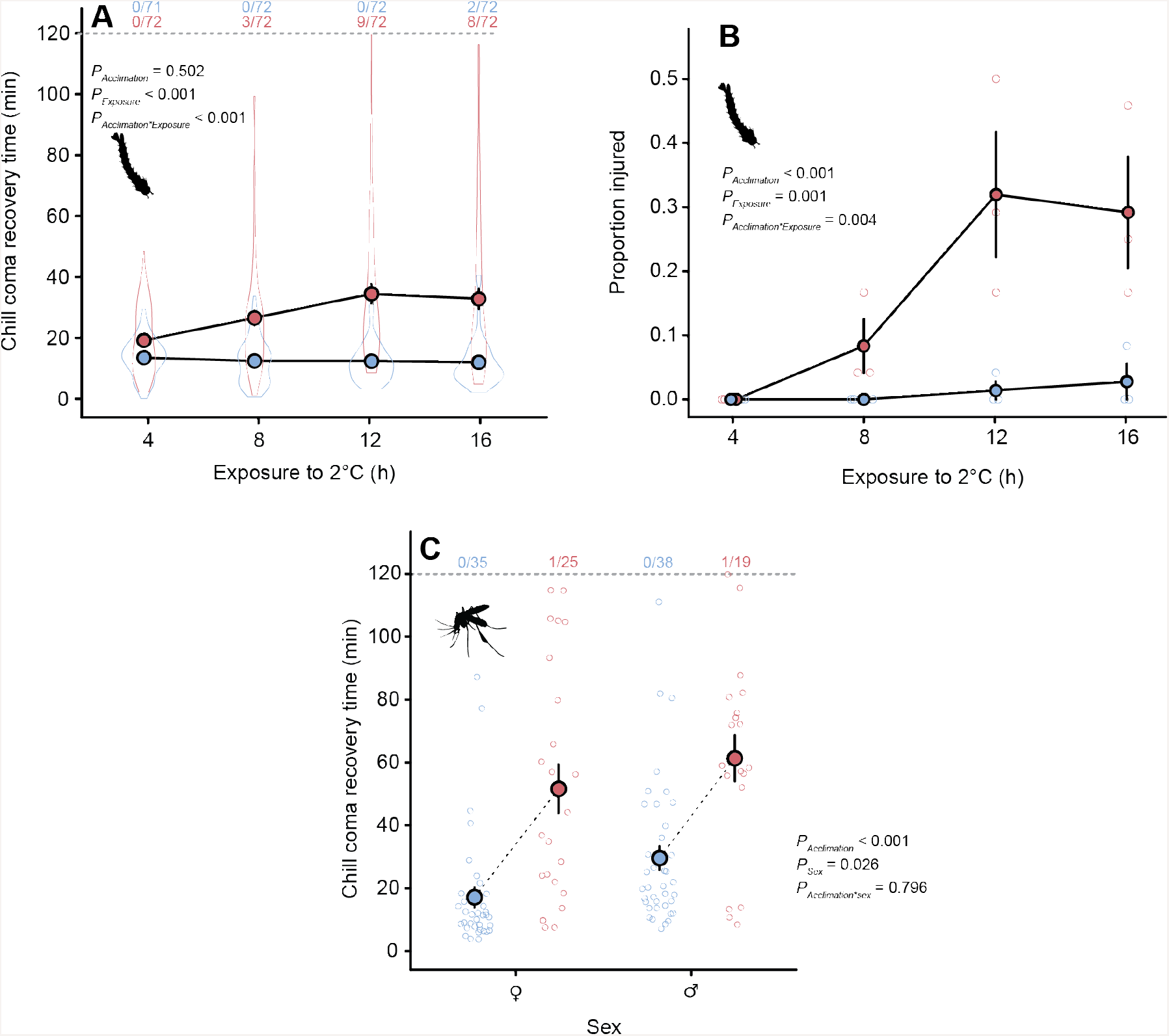
Chill coma recovery times (CCRT) of larval (A) and adult (B) warm- (25°C) and cold (15°C) *Aedes aegypti*. Larvae were exposed to 2°C for one of four different durations (x-axis) and adult CCRT was recorded following 6 h at 2°C. Violin plots in panel A represent sample distributions (owing to large sample size). Open circles in panel B represent proportion of mosquitoes injured (unable to resume siphon use within 2 h) in three independent trials. Open circles in panel C represent individual adult mosquitoes. Ratios above the dashed lines in panels A and C represent the number of individuals that did not recover from chill coma within the observation period (120 min). In all panels closed circles represent the mean (± sem) of each group. Error bars that are not visible are obscured by the symbols.

Cold acclimation also significantly improved rates of chill coma recovery in adult mosquitoes; Cold-acclimated mosquitoes recovered from chill coma after 6 h at 0°C approximately 25 min faster than the warm-acclimated conspecifics (Fig. 2C; main effect of acclimation: F_3,112_ = 38.1, *P* < 0.001). As with CCO, females appeared more cold tolerant based on CCRT, and recovered ~10 min faster than males (on average) from the same cold stress (main effect of sex: F_3,112_ = 5.1, *P* = 0.026). Unlike CCO, there was no interactive effect of sex and acclimation temperature on chill coma recovery (F_3,112_ = 0.01, *P* = 0.796), as cold acclimation improved adult CCRT by the same degree (~32-34 min) regardless of sex (Fig. 2C).

Warm-acclimated mosquitos were far more likely to take a blood meal when it was offered (Fig. 3A; t = 26.3, *P* < 0.001); only two cold-acclimated females (out of 65 that were offered it) voluntarily fed on blood within 20 min (Fig. 3A). For cold-acclimated mosquitos that did not feed, exposure to a blood meal still impacted CCRT, as increasing time since exposure to the blood led to longer recovery times (Fig. 3B; F_1,61_ = 4.1, *P* = 0.046). For warm-acclimated mosquitos, the act of blood feeding had no effect on CCRT (Fig. 3C; main effect of feeding status: F_1,61_ = 0.2, *P* = 0.889), and although there was a slight tendency for CCRT to increase with time since the blood was offered, this effect was not statistically significant (main effect of time: F_1,61_ = 2.8, *P* = 0.100), and there was no significant interactive effect of feeding status and time on CCRT (Fig. 3C; F_1,61_ = 0.1, *P* = 0.838).

**Figure 3:**
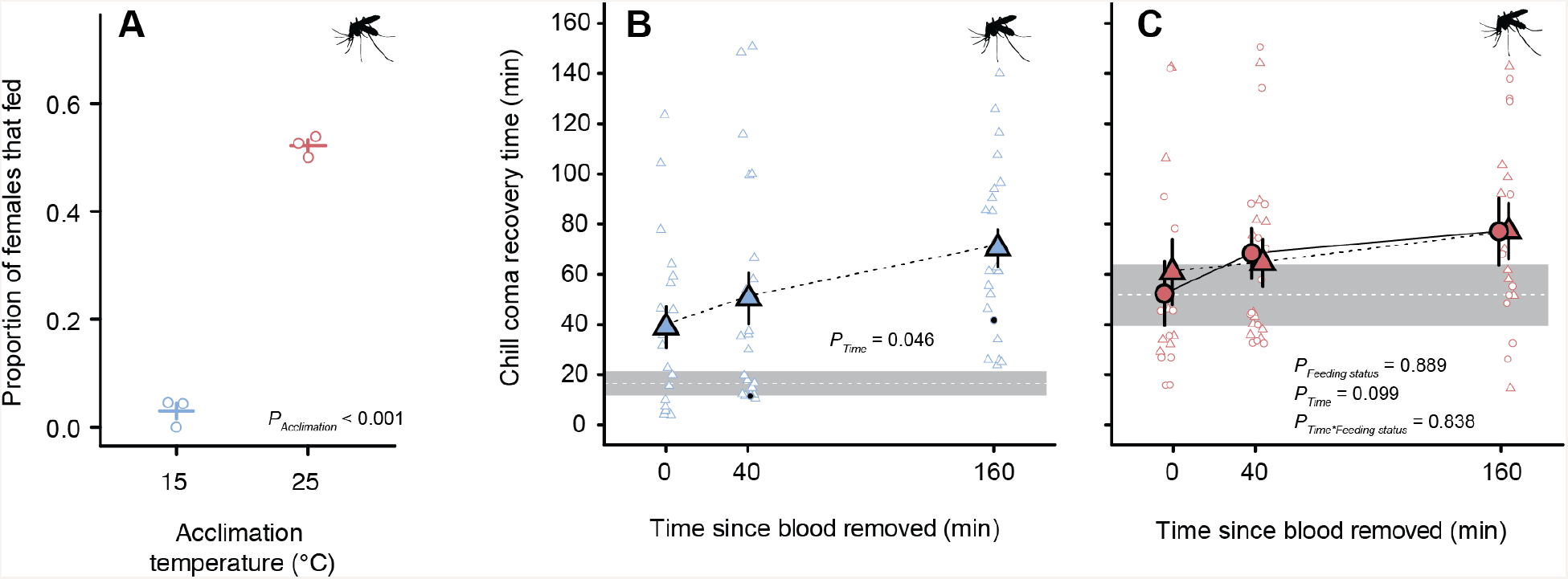
Rates of blood feeding (A) and chill coma recovery times following blood feeding of cold- (B; 15°C; blue) and warm-acclimated (C; 25°C, red) adult *Aedes aegypti* females. Mosquitoes of both acclimation groups were offered warm sheep’s blood for 20 min before the blood was removed. Individual mosquitoes were then given 0 min or an additional 40 or 160 min at room temperature before they were exposed to 0°C for 6 h. Triangles represent mosquitoes that chose not to feed on the blood while circles represent those that did feed. Open circles in panel A represent the proportion of mosquitoes that fed within each treatment group. Open symbols in panels B and C represent CCRT values of individual adult mosquitoes. The two closed circles in panel B represent the two cold-acclimated mosquitos that took a bloodmeal (see Results text). In all cases, closed blue and red symbols represent the mean (± sem). Error bars that are not visible are obscured by the symbols.

### Low temperature survival

The most parsimonious model for larval survival retained the interaction between exposure temperature and acclimation temperature, which significantly interacted to determine survival (z = 2.6, *P* = 0.010). Temperature strongly influenced larval survival in both acclimation groups (main effect of temperature: z = 9.4, *P* < 0.001), with larvae exposed to lower temperatures suffering higher mortality (Fig. 4A). Cold-acclimated larvae survived 24 h exposures to lower water temperatures (LT_50_ = −1.64 ± 0.23°C) than their warm-acclimated conspecifics (LT_50_ = 0.81 ± 0.18°C; main effect of acclimation temperature: z = 7.49, *P* < 0.001; Fig. 4A). The most parsimonious model of adult survival at low temperatures eliminated all interactions between acclimation temperature, exposure temperature, and sex, but retained all of these variables as independent effects. Adult mosquitoes exposed to lower temperatures suffered greater mortality (main effect of exposure temperature: z = 12.0, *P* < 0.001), and as was the case for both chill coma onset and chill coma recovery, adult female mosquitoes were consistently more cold-tolerant than males (Fig. 4B; main effect of sex z = 2.4, *P* = 0.014). For both sexes, cold-acclimated adults survived to lower temperatures than warm-acclimated adults (Fig. 4B; main effect of acclimation temperature: z=7.1, *P* < 0.001). Cold acclimation shifted the female LT_50_ (following 6 h cold exposures) from 0.0 ± 0.19°C to −1.9 ± 0.15°C and the male LT_50_ from 0.3 ± 0.18°C to −1.3 ± 0.19°C (Fig. 4B).

**Figure 4:**
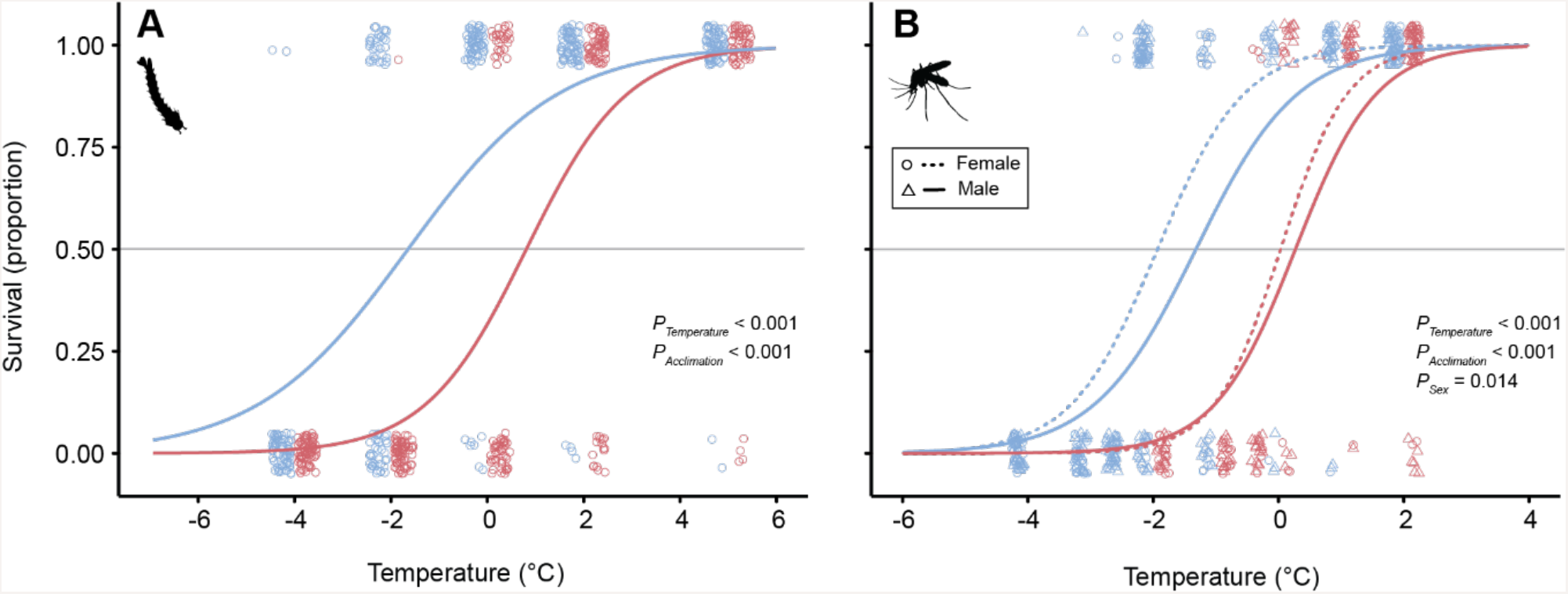
Rates of survival following cold exposure in larval (A) and adult (B) warm- (25°C; red) and cold-acclimated (15°C; blue) *Aedes aegypti*. Open symbols represent individual mosquitoes and are slightly shifted (both vertically and horizontally) for visual clarity. Lines represent models of best fit. Larvae were exposed to treatment temperatures for 24 h and adults for 6 h.

### Hemolymph ion balance

Exposure to 0°C for 24 h caused both warm- and cold-acclimated larvae to lose hemolymph [Na^+^] balance. Both acclimation groups had similar hemolymph [Na^+^] prior to cold stress, and cold stress caused hemolymph [Na^+^] to significantly decrease in both groups (Fig. 5A; main effect of cold exposure: F_3,83_ = 74.1, *P* < 0.001). There was no main effect of acclimation treatment on [Na^+^] (F_3,83_ = 0.1, *P* = 0.800), nor any interaction between acclimation treatment and cold exposure (F_3,83_ = 0.1, *P* = 0.731). Cold exposure also caused both warm- and cold-acclimated larvae to lose hemolymph K^+^ balance. Cold stress elevated hemolymph [K^+^] in both groups (Fig. 5B; main effect of cold exposure: F_3,84_ = 43.4, *P* < 0.001). Notably, this effect of chilling on hemolymph [K^+^] was more pronounced in warm-acclimated than cold-acclimated larvae (Fig. 5B; interaction between acclimation group and cold exposure: F_3,84_ = 6.4, *P* = 0.013); 24 h at 0°C cold exposure elevated mean hemolymph [K^+^] in warm-acclimated mosquitos by ~130%, but only by 65% in cold-acclimated larvae (Fig. 5B). Because of this difference following cold stress (and because cold-acclimated larvae tended to have very slightly lower mean hemolymph [K^+^] prior to cold stress: 4.1 mM vs. 4.3 mM) there was also a significant main effect of acclimation group on hemolymph [K^+^] (F_3,84_ = 6.8, *P* = 0.011).

**Figure 5:**
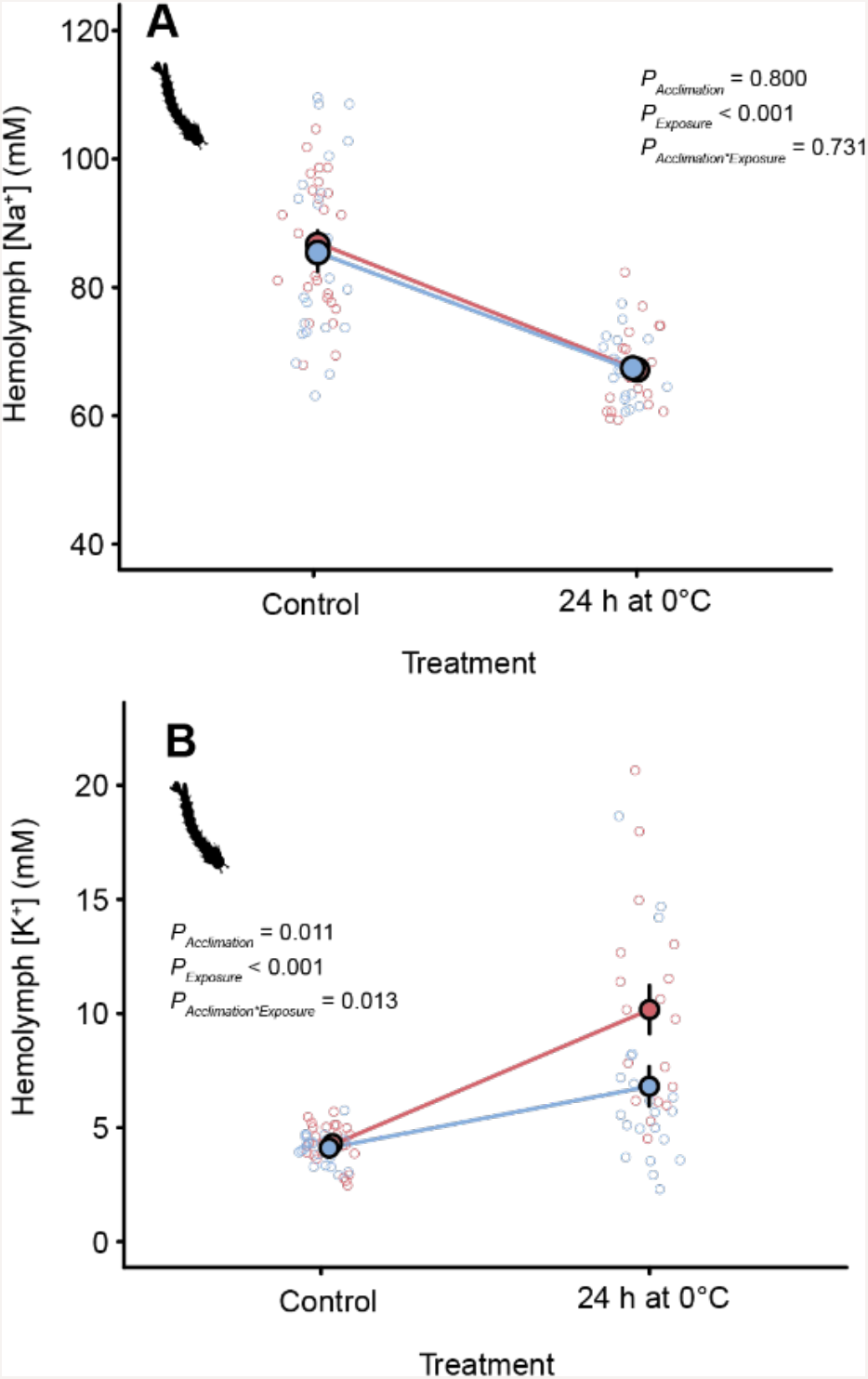
Concentrations of Na^+^ (A) and K^+^ (B) before and after cold stress in larval *Aedes aegypti*. Open circles represent individual samples and closed circles represent the mean (± sem). Error bars that are not visible are obscured by the symbols.

## Discussion

Larvae and adults of *Aedes aegypti* are clearly capable of cold acclimation when presented with a change in developmental or adult acclimation temperature. In the present study we compared the effects of development or adult acclimation at only two temperatures (15°C and 25°C), but demonstrate that this difference of 10°C was sufficient to substantially alter chilling tolerance in this important vector of disease. Cold-acclimated larvae and adults more rapidly recovered from chill coma following cold stress, and had significantly higher survival following chronic cold. After 12-16 h at 2°C, very few larvae acclimated to 15°C showed any signs of chilling injury while ~30% of larvae acclimated to 25°C were clearly suffering from neuromuscular injury that prevented them moving in a coordinated manner (Fig. 2B).

Chilling injury has been repeatedly associated with a systemic loss of ion balance in several terrestrial insects, including members of Hemiptera, Diptera, Blattodea, Lepidoptera, and Orthoptera (M. K. Andersen, Jensen, & Overgaard, 2017; V Koštál et al., 2006; Vladimír Koštál, Vambera, & Bastl, 2004; MacMillan & Sinclair, 2011b; MacMillan, Andersen, Davies, et al., 2015; MacMillan, Findsen, Pedersen, & Overgaard, 2014). Notably, however, all tests of the ionoregulatory collapse model have been previously done on terrestrial insects. Here, we demonstrate a similar inability to maintain low hemolymph [K^+^] in the cold in an aquatic larval insect (Fig. 5). In several terrestrial insects, cold acclimation improves chilling tolerance and prevents hyperkalemia. In such cases, hyperkalemia is mitigated (at least in part) through modifications to renal ion and water transport that help to clear excess K^+^ ions from the hemolymph and maintain hemolymph volume (M. K. Andersen, Folkersen, et al., 2017; MacMillan, Andersen, Loeschcke, et al., 2015; Yerushalmi, Misyura, MacMillan, & Donini, 2018). Although at present it is unclear whether the same mechanisms underlie improvements in chilling tolerance in mosquito larvae, the prevention of hyperkalemia likely attenuates cold-induced cell membrane depolarization, which would limit cell death and thereby facilitate survival (M. K. Andersen, Folkersen, et al., 2017; Bayley et al., 2018; Boutilier, 2001; MacMillan, Baatrup, et al., 2015).

We noted that the majority of *Aedes* larvae tend to sink upon entering chill coma. Mosquito larvae obtain gaseous oxygen from the water surface through a siphon on the posterior end of their abdomen, so sinking during cold stress may limit access to oxygen during a cold stress and cause systemic hypoxia. Like cold stress, anoxia has been demonstrated to cause disruptions of ion homeostasis leading to hyperkalemia in *Drosophila* (Campbell, Andersen, Overgaard, & Harrison, 2018), meaning an inability to access sufficient oxygen during chill coma may further contribute to ionic imbalance and injury in the cold in this aquatic insect. Alternatively, as the metabolic rate of ectotherms is strongly supressed during cold exposure, larvae may obtain sufficient oxygen from the surrounding water during cold stress to fuel metabolism and avoid the downstream consequences of hypoxia.

We were surprised to find that cold-acclimated adult mosquitos displayed a very strong aversion to blood feeding when the opportunity was presented (Fig. 3). As a tropical and subtropical species, *Aedes aegypti* is not known to be capable of any form of quiescence or diapause (Diniz et al., 2017), but a reduction in feeding behaviour is one of several hallmarks of insects in a period of dormancy, including mosquitos. We will not speculate on whether or not some manner of dormancy is taking place in cold-acclimated *Aedes aegypti* but argue that this subject is worthy of further investigation, particularly given the importance of this species to human health. Despite not feeding, chill coma recovery times of female mosquitos that were simply in the presence of blood increased (became worse) over the 3 h following its presentation. Most likely, this reduction in cold tolerance was driven by the warmth of the blood, which may induce rapid changes in thermal tolerance in exposed mosquitos. Drinking a blood meal induces an adaptive heat shock response in *Ae. aegypti* that protects against the effects of a rise in body temperature on fecundity (Benoit et al., 2011). We thus hypothesize that either the temperature of the warm blood or some other signal of its presence induces a similar response that alters mosquito thermal tolerance. In contrast to cold-acclimated mosquitoes, approximately half of the warm-acclimated females fed on blood within 20 mins of its presentation (Fig. 3). Although we hypothesized that the salt load associated with a blood meal would alter ionoregulatory homeostasis and thereby alter chill tolerance, there was no effect of blood feeding on CCRT in warm-acclimated mosquitos. There was a tendency for the CCRT of warm-acclimated mosquitos to increase over time following presentation of the blood (as was seen in cold-acclimated adults), this trend was not statistically significant, possibly because the acclimation temperature (25°C) was closer to the temperature of the blood.

The adult chill coma onset temperature (CCO) of *Ae. aegypti* appears highly plastic (Fig. 1B), as cold acclimation reduced the CCO of female and male mosquitoes by approximately 6.4 and 3°C, respectively. In stark contrast to adults, however, larvae acclimated to 15°C had the same CCO as those acclimated to 25°C (~6°C; Fig. 1A), despite being more tolerant of chilling by every other measure. The CCO, CCRT, and chilling injury are all thought to be related to the capacity to maintain ion and water balance, but are mediated by different specific physiological mechanisms of failure occurring in different organs and across different time scales (MacMillan, 2019; Overgaard & MacMillan, 2017; Robertson et al., 2017). Our results in the present study thus suggest that acclimation alters mechanisms underlying CCRT and the development of chilling injury without impacting the temperature that causes paralysis. Further, this result suggests that for larvae, measuring the CT_min_ or CCO alone may strongly underestimate variation in cold tolerance in this species. We thus strongly recommend that other measures of cold tolerance (e.g. survival following cold stress) be included in future comparisons of thermal tolerance among or within populations, particularly in the study of larval thermal tolerance. Winter temperatures appear to be a critically important predictor of the suitability of environments for the persistence of *Ae. aegypti* and *Ae. albopictus* (Johnson et al., 2017) and both species are spreading into habitats that have been previously considered too cold for their permanent establishment. Recent studies in *Drosophila* and other insects have suggested that range limits are closely associated with the frequency and severity of temperatures crossing critical physiological thresholds that mark the boundaries of activity (e.g. CT_min_, CCO) or survival (J. L. Andersen et al., 2015; Bozinovic, Calosi, & Spicer, 2011; Calosi, Bilton, Spicer, Votier, & Atfield, 2010; Overgaard et al., 2014). Although typically considered a tropical species with little capacity for overwintering, *Ae. aegypti* appears to have a substantial thermal acclimation capacity, and this ability is at least partly associated with an improved ability to prevent cold-induced hyperkalemia. Given that acclimation can alter thermal limits, thermal plasticity is likely to be an important factor governing the ability for invasive species like *Ae. aegypti* to survive in new environments and respond to the effects of climate change on the mean and variance of environmental temperatures. The experimental population used in the present study is derived from strains held in a laboratory environment for decades, and acclimation capacity can vary widely in insects in the wild. Eggs of *Ae. aegypti* at the southern end of their American range appear to have evolved greater cold tolerance than populations previously studied (De Majo et al., 2017), so a careful analysis of population-level variation in thermal tolerance plasticity in *Ae. aegypti* is overdue, and would serve to inform future models of the distribution of this dangerous disease vector.

## Supporting information

Data archive

## Data Accessibility

All data is provided as a supplementary file for review and the same file will be uploaded to a data repository should the manuscript be accepted for publication.

## Author Contributions

All authors contributed to the conception and design of the study, A.J. H.D. and G.Y. conducted the experiments. A.J., G.Y., H.D. and H.M. analyzed the data, H.M. drafted the manuscript, and all authors edited the manuscript.

